# The creation and selection of mutations resistant to a gene drive over multiple generations in the malaria mosquito

**DOI:** 10.1101/149005

**Authors:** Andrew Hammond, Kyros Kyrou, Marco Bruttini, Ace North, Roberto Galizi, Xenia Karlsson, Francesco M Carpi, Romina D’Aurizio, Andrea Crisanti, Tony Nolan

## Abstract

Gene drives have enormous potential for the control of insect populations of medical and agricultural relevance. By preferentially biasing their own inheritance gene drives can rapidly introduce genetic traits even if these confer a negative fitness on the population.

We have recently developed gene drives based on a CRISPR nuclease construct that is designed to disrupt key genes essential for female fertility in the malaria mosquito. The construct copies itself and the associated genetic disruption from one homologous chromosome to another during gamete formation, in a process called homing that ensures the majority of offspring inherit the drive. Such drives have the potential to cause long-lasting, sustainable population suppression though they are also expected to impose a large selection pressure for resistance in the mosquito. One of these population suppression gene drives showed rapid invasion of a caged population over 4 generations, establishing proof of principle for this technology. In order to assess the potential for the emergence of resistance to the gene drive in this population we allowed it to run for 25 generations and monitored the frequency of the gene drive over time. Following the initial increase of the gene drive we noticed a gradual decrease in its frequency that was accompanied emergence of small, nuclease-induced mutations at the target gene that are resistant to further cleavage and restore its functionality. Such mutations show rates of increase consistent with positive selection in the face of the gene drive. Our findings represent the first documented example of selection for resistance to a synthetic gene drive and lead to important design recommendations and considerations in order to mitigate for resistance for future gene drive applications.

## INTRODUCTION

Naturally occurring gene drives – selfish genetic elements that are able to bias their own inheritance and rapidly invade a population, even starting from very low frequencies – have inspired proposals to harness their power to spread into a population of insect disease vectors traits that manipulate the biology of insect vectors of disease in ways that could suppress or eliminate disease transmission^1-3^. In particular for malaria, transmitted exclusively by mosquitoes of the Anopheles genus, historical gains in reducing the disease burden have been largely achieved by the correct implementation of vector control measures (residual insecticides and bed nets) ^4^. Though these measures have been instrumental in substantially reducing malaria transmission they are deemed unsuitable to eradicate the disease in the near future at the current level of investment^5^. Gene drive technology could help in developing a self-sustaining, species-specific and affordable vector control measure much needed to achieve disease eradication in the future. Gene drives based on the activity of DNA nucleases able to recognise very specific target sequences were first proposed over a decade ago and have received much attention recently due to the advent of new, easily programmable nucleases such as CRISPR-Cas9 that have allowed us and others to build functioning gene drives that show rates of inheritance close to 100%, compared to the expected Mendelian inheritance of 50% ^1, 6-8^. The principle behind the technology is to re-program a nuclease to cleave a specific site of interest in the genome and to insert the nuclease within this recognition site. The gene drive is designed to be active in the germline, so that in diploid organisms heterozygous for the gene drive the nuclease causes a double stranded break (DSB) at the target site on the homologous chromosome not containing the gene drive. The DSB can be repaired either by simple end-joining (EJ) of the broken strands or via homology-directed repair (HDR) where the DSB is resected and the intact chromosome used as a template to synthesise the intervening sequence. In the case of a gene drive nuclease repair via HDR thus leads to a copying of the drive from one chromosome to another and the conversion of a heterozygote into a homozygote. Hence the force of gene drive is determined by a combination of the rate of cleavage of the nuclease in the germline, and the propensity for the cell machinery to repair the broken chromosome by HDR. We and others have shown that in germline cells the rates of HDR following a nuclease-induced DSB can be almost two orders of magnitude greater in the germline than EJ, a fact which explains the extraordinarily high rates of gene drive inheritance observed. On the other hand EJ repair can lead to the creation of small insertions or deletions at the target site that, although occurring initially at low frequency, might be expected to be selected for in the target organism if they prevent the gene drive nuclease acting and there is a negative fitness cost associated with the gene drive ^1, 6, 9-11^. This possibility has been recognised since the first proposal of this type of gene drive ^1^with much theory being dedicated to it recently ^11, 12^. There are several potential mitigation strategies including, but not limited to, the targeting of conserved sequences that are less tolerant of mutations and the targeting of multiple sequences, akin to combination therapy, in order to lower the likelihood of resistance arising ^1, 9^. We previously developed a gene drive designed to spread into a mosquito population and at the same time reduce its reproductive potential by disrupting a gene essential for female fertility, thus imposing a strong fitness load on the population. To investigate the long term dynamics of the emergence of resistance to a gene drive imposing such a load we continued to monitor the frequency of this gene drive over generations and analysed the target locus for evidence of mutagenic activity that could lead to the development of resistant alleles that block gene drive activity and restore gene functionality. Our findings show that a range of different resistant alleles can be generated with differing frequencies and some of these are subsequently selected for and show dynamics consistent with our modelling predictions. These results provide a quantitative framework for understanding the dynamics of resistance in a multi-generational setting and allow us to make concrete recommendations for the improvement of future gene drive constructs that relate to choice of target site and regulation of nuclease expression in order to reduce the emergence of resistance.

## RESULTS

A proof-of-principle CRISPR-based gene drive designed for population suppression was previously developed in our laboratory (Fig 1A). This gene drive disrupted a haplosufficient gene (AGAP007280, the putative mosquito ortholog of *nudel^13^)* required in the soma and essential for female fertility ^10^. The gene drive also contained an RFP marker gene for the visual detection of individuals inheriting the drive. In our experiments individuals heterozygous for the gene drive transmitted the drive, regardless of their sex, to more than 99% of their offspring. We observed in these mosquitoes a marked reduction in fertility (~90%) in females heterozygous for the drive, due to ectopic expression of the nuclease under control of the germline vasa2 promoter that resulted in conversion to the null phenotype in somatic cells. In spite of this fitness disadvantage both our modelling and experimental data showed that the gene drive could increase rapidly in frequency in a caged population due to the exceptionally high rates of inheritance bias. From a starting population (G_0_) in two duplicate cages of 600 with a 1:1 ratio of transgenic heterozygotes and wild type individuals, the gene drive progressively increased in frequency to 72-76.5% by G_4_. This rate of increase was slightly higher than predicted by a deterministic model but within the limits of stochastic variation expected. Due to a combination of the partial dominance of the sterility phenotype in heterozygous females and the previously documented generation of target site mutations conferring resistance to the gene drive ^10^, this first gene drive was not expected to maintain high levels of invasion. Nonetheless it represented a useful experimental model to investigate the long term dynamics of the de novo generation of target site mutations and their selection at the expense of a gene drive imposing a large reproductive load. We therefore maintained this cage experiment for 25 generations and used the presence of the RFP marker in the gene drive construct as a proxy to estimate the frequency of individuals containing it. The frequency of gene drive progressively increased in both duplicated cages peaking at around generation 6, thereafter we observed a gradual and continuing decrease such that by G_25_ the frequency of individuals with the gene drive was less that 20%. To investigate whether the drop in the gene drive frequency was due to the selection of pre-existing variant target sites in the population or the generation and selection of nuclease-resistant indels we used deep sequencing of a PCR amplicon comprising sequences flanking the target site on pooled samples of mosquitoes from early (G_2_) and late (G_12_) generational time points (Fig 1B). The expected amplified region from the original wild type sequence was 320bp long, with the putative cleavage point within the target site residing after nucleotide 203 (Fig 2A). Ultra-deep sequencing of PCR reactions performed on pooled DNA under non-saturating conditions so that the number of reads corresponding to a particular allele at the target site is proportional to its representation in the pool. In the colony of mosquitoes that we used there are a number of pre-existing single nucleotide polymorphisms (SNPs) within the amplicon that do not overlap the nuclease target site and are present at varying frequencies. We developed a computational method to analyse the sequences edited by the CRISPR-based gene drive close to the nuclease target site. The method was specifically designed to identify small insertions and/or deletions eventually introduced by the repair system and also to characterize the haplotypes on which they arise. Because the PCR amplicon only represents the non-drive allele the frequencies reported refer to their frequency within this class, rather than within the population as a whole. Mapping amplicon sequences reconstructed from the sequenced against the Anopheles gambiae reference genome (PEST strain, AgamP4, Vectorbase) already in the G_2_ generation we noticed a vast repertoire of deletions, with a wide range of sizes and centred around the predicted nuclease cleavage site after nucleotide 208 (Fig 2A - one cage trial shown as a representative example), with a relatively lower proportion of small insertions, consistent with the known mutational activity of the nuclease. On the contrary ten generations later we observed an obvious stratification of deletions and a much narrower distribution. To investigate whether the change in distribution reflected a selective advantage for specific indel alleles in the population, rather than a biased propensity for their de novo generation, we examined the nature of each allele and its frequency over time (Fig 2B). We considered all alleles that reached a frequency of at least 1*%* in any condition and classified these as to whether the indel caused a frameshift in the coding sequence of the target gene or was in-frame. The predominant target allele in the G2 was still the reference (non-mutated) allele at 63 and 48% in replicates 1 and 2, respectively (Fig 2B and Supp Table 1), while the second major class (at least 15% in each replicate) was represented by a wide range of non-reference alleles, each present at low frequency (<1%) consistent with the stochastic generation of a broad range of indels. Thus at a time when the gene drive was still increasing in frequency in the population there was a significant silent accumulation of mutations at the target site that would likely render it refractory to the homing mechanism of copying, Notably, three separate indels causing in-frame deletions of 3- or 6bp (202-TGAGGA, 203-GAGGAG, 203-GAG; where 203 refers to the starting site of the indel in the reference amplicon and "-" means deletion) were present at low but appreciable frequencies in the G_2_. Such short in-frame deletions may result in only minimal disruption to the final encoded protein while at the same time proving resistant to the nuclease component of the gene drive. Indeed these three deletions, plus a 6bp in frame insertion (207+AAAGTC) had increased significantly in frequency to make up the 4 most abundant non-drive alleles in the G_12_, almost to the exclusion of the reference allele (present at 6% and 0.4%; supp table 1). Notably a wide range of frameshift indels that were present in G_2_ had fallen in frequency in G_12_ to either below the 1% threshold or were not detected at all (Fig 2B and Supp Table 1). The most parsimonious explanation for these trends is that the in-frame indels are being selected for because they restore functionality to the target gene while protecting the sequence from gene drive activity that otherwise would impair female fertility. These ‘restorative’ mutations are likely to be most strongly selected when the frequency of the gene drive is very high in the population – when the majority of individuals are homozygous the relative gain in viable offspring from an individual with a gene drive balanced by a resistant restorative mutation is that much higher. In theory small in-frame deletions could arise by either classical non-homologous end joining (NHEJ), or an alternative form of end-joining (microhomology-mediated end joining, MMEJ) that relies on alignment of small regions of microhomology, as little as 2 base pairs, on either side of the DSB resulting in loss of the intervening sequence ^14^. In agreement with this possibility at least three of the most frequent alleles in the G_12_ generation can feasibly be explained by MMEJ via 3bp repeats (Fig 2C). We used the presence of naturally occurring SNPs to investigate the haplotypes associated with those indels at highest frequency after G_12_. This analysis showed that in cage 1 the deletion 203-GAGGAG was present on 10 separate haplotypes in G_12_, with each haplotype being present at ratios broadly similar to their ratios in the starting population (Supp Fig 1), suggesting that the same deletion was generated at least 10 times independently and that there was no obvious selective advantage to any particular haplotype surrounding the deletion. In cage 2 the predominant allele 202-TGAGGA (68% of all non-reference alleles), an end-joining deletion that shows no apparent features of a MMEJ event, was formed on 5 separate haplotypes. The progressive increase in frequency of these mutations at the target site, concomitant with a decrease of gene drive activity, strongly suggested that they conferred resistance to cleavage while still ensuring a normal functional activity of *nudel* thereby explaining their selective advantage. To confirm this hypothesis we crossed individual RFP+ females from G_20_ with wild type males and assessed both their fertility and the transmission rate of the drive. We also sequenced the target site of each parental female to characterize allelic variants at the target locus. This analysis failed to detect wild type sequence at the target site (Fig 3A) among 70 individuals tested, instead every individual showed an indel indicating that each female tested was heterozygous for the gene drive and balanced by a mutated target site; in replicate CT1 the 203-GAGGAG 6bp deletion was the predominant allele (23/31 individuals) while in CT2 another 6bp deletion (202-TGAGGA) was predominant (37/39 individuals). The relative frequency of each allele was consistent with the results obtained using pooled amplicon sequencing performed on the G12 individuals. Importantly, of these heterozygous females that could be confirmed as mated the vast majority (56/58) generated viable progeny (average clutch size 119 +/- 35.9 eggs, average hatching rate 78.5% +/-19.9%; Fig 3B and Supp Table 2) at rates significantly higher than those previously observed when crossing the gene drive in heterozygosity with a wild type target site (90.7% overall reduction in fecundity)^10^ thus suggesting that the mutations detected at the target site did not impair the functionality of *nudel*.

**Figure 1.**
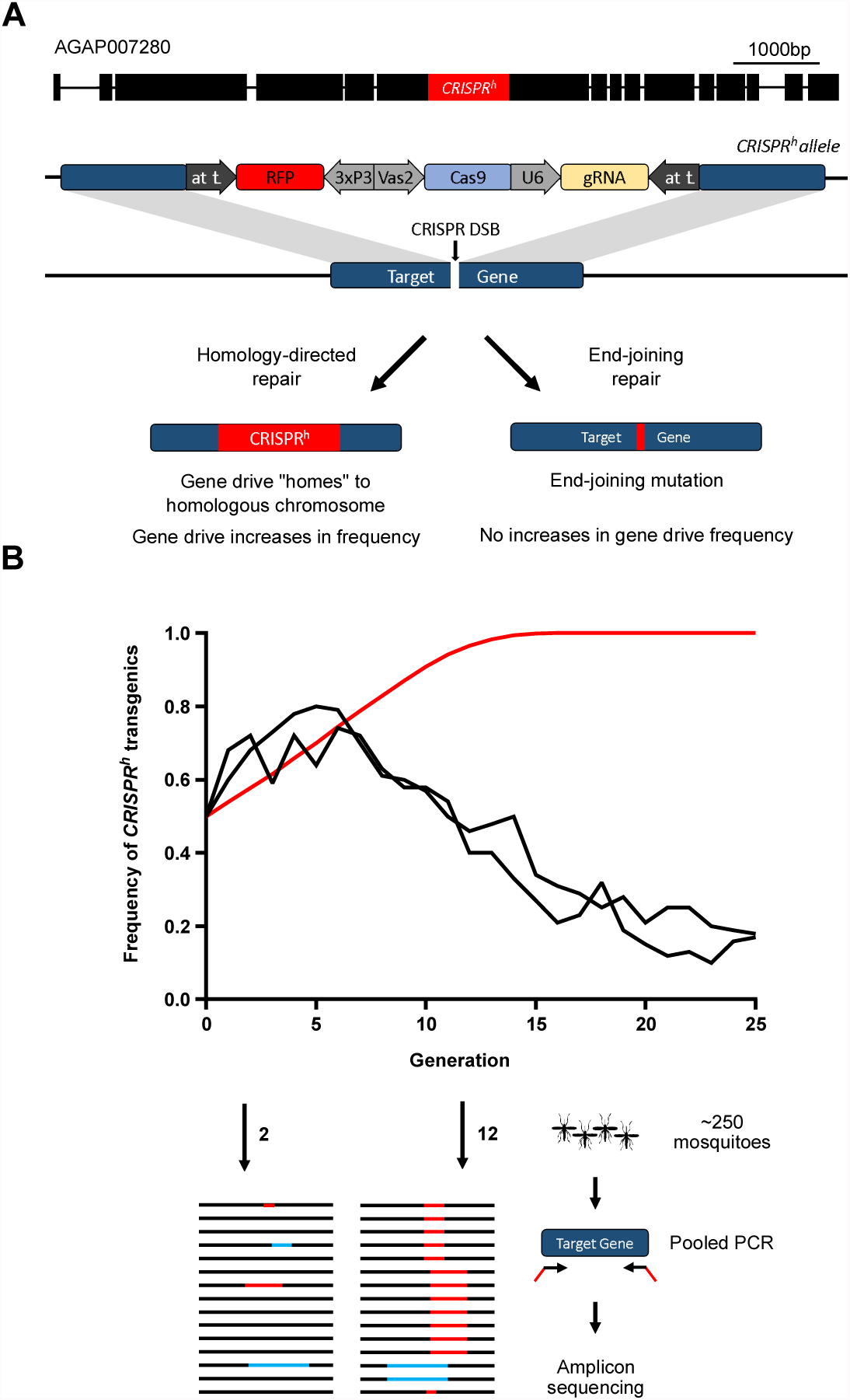
Dynamics of a population suppression gene drive construct over 25 generations. **(A)** Design of the CRISPR-based gene drive construct and the relevant position of its target site within AGAP007280; **(B)** The proportion of individuals containing at least one copy of the gene drive in two replicate cages, monitored each generation for 25 generations. Black lines represent the observed frequencies in each of the two cage trials (CT1 and CT2), red line represents the predicted frequency according to the previous deterministic model that did not take into account target site resistance (Hammond et al.). Samples were taken for pooled sequencing analysis of the gene drive target site at G_2_ and at G_12_.

**Figure 2.**
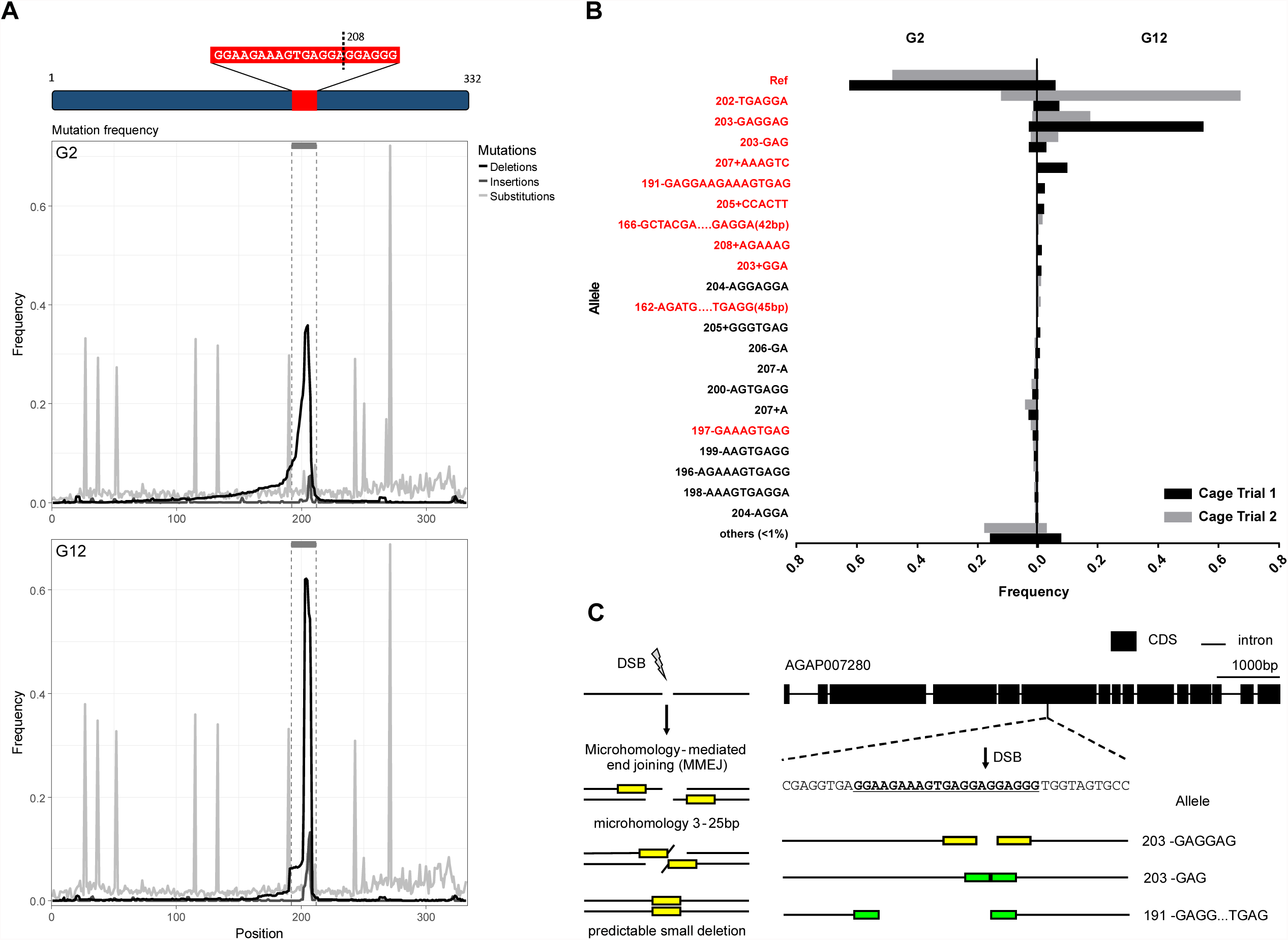
Nuclease-generated target site mutations show selection over time. **(A)** Location of the target site of the CRISPR-based gene drive relative to the region amplified for pooled sequencing (highlighted in red), with the double stranded break occurring after nucleotide 208. Cumulative frequencies of deletions (black), insertions (dark grey) and substitutions (light grey) at each nucleotide position along the amplicon were plotted. Numerous substitutions outside of the target site represent SNPs circulating in the lab strain of mosquito in which the gene drive was introduced and were used for analysis of haplotypes of target site mutations (Supp Fig. 1). One cage trial (CT1) shown as a representative example. **(B)** Frequency of individual target site alleles in both G_2_ and G_12_ for cage 1 (black) and cage 2 (grey) with in-frame indels highlighted in red. All alleles never exceeding 1% frequency in at least one condition were grouped together as one class (<1%). **(C)** Microhomologies flanking the double stranded break at the target could explain some of the most frequent deletions observed. 3bp microhomologies of AGG (yellow) or GGA (green) and the resultant deletion alleles arising in the event of MMEJ are shown.

**Figure 3.**
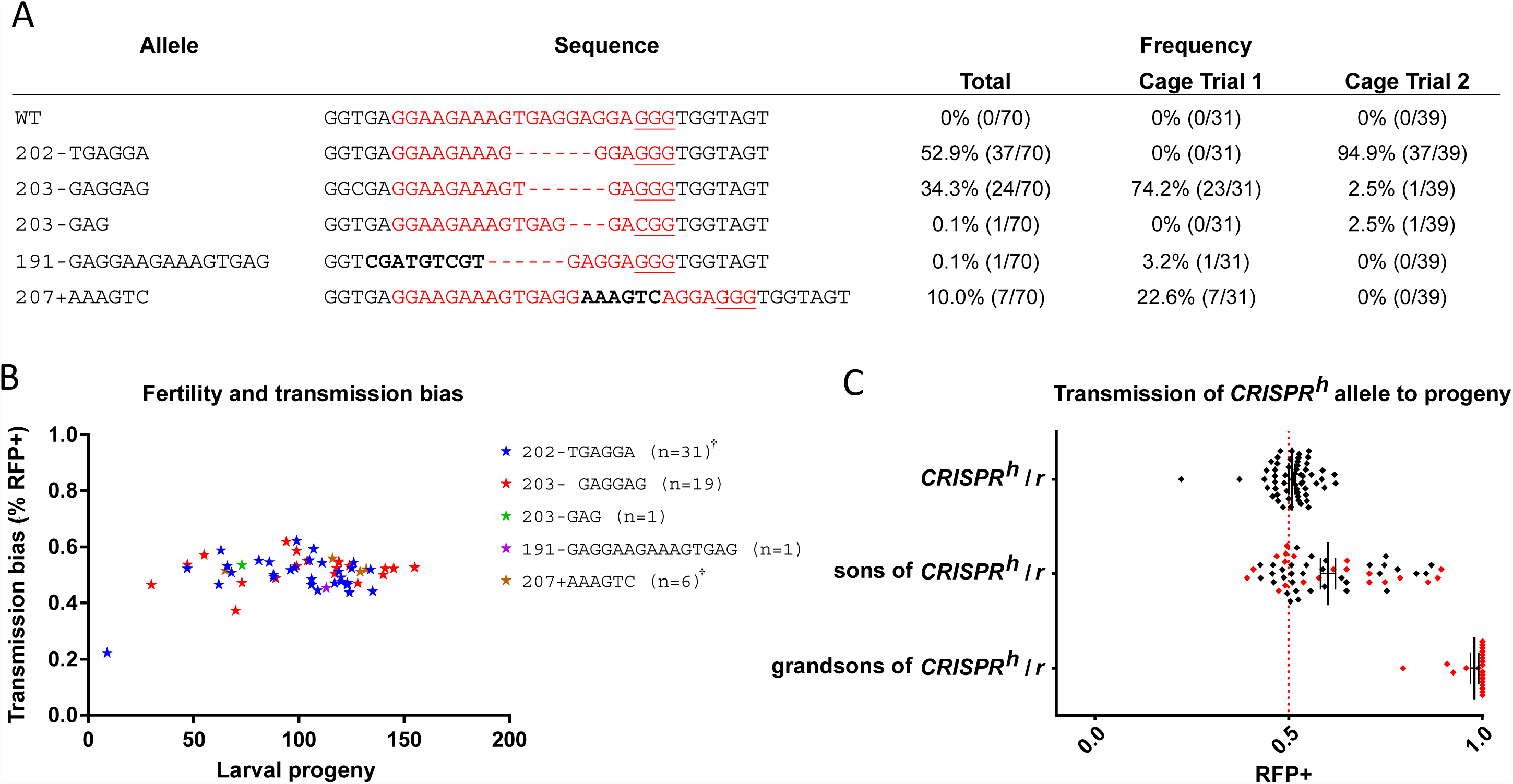
Target site mutations under positive selection are resistant to gene drive activity and restore function to the target female fertility gene. **(A)** Individual females containing at least one copy of the gene drive (RFP+) were selected from the G_20_ generation and the nature of the target allele was determined by PCR and sequencing. Each class of allele is shown with gene drive target sequence highlighted in red and PAM sequence underlined. **(B)** The fecundity of these females and transmission rates of the gene drive were measured and grouped according to allele class at the target site. **(C)** Each *CRISPR^h^/r* female was used to form a separate lineage and transmission of the gene drive was assessed in sons receiving a maternal copy of the gene drive. A smaller fraction of grandsons receiving a paternal copy of the gene drive were similarly assessed for gene drive transmission. Individual lineages assessed in all three generations are marked in red. † Of 58 mated females one (with deletion 207+AAAGTC) failed to produce eggs while another (202-TGAGGA) produced eggs that failed to hatch.

The offspring of these individuals showed a non-biased inheritance of the gene drive (average inheritance 50.82% +/-5.96%, total RFP+ offspring 50.85%) consistent with normal Mendelian segregation of the element (Fig 3B) thus indicating that the mutated sequences were resistant to nuclease activity. There was no obvious difference in fertility associated with the presence of the 5 classes of indel sampled in this assay (Fig 3B). The two major indels across the two cages result in relatively conservative changes to the overall amino acid sequence of the final gene product - two glutamate residues missing (203-GAGGAG) or two glutamates missing and a conversion of a lysine to arginine (202-TGAGGA) - in the final protein product.

Conceivably a breakdown in the nuclease component (e.g. mutation in the Cas9 coding sequence, or a mutation in the gRNA sequence) could be an alternative explanation for the Mendelian transmission of the RFP-marked gene drive element and restored fertility in heterozygous females that we saw. To assess this possibility we took the male offspring (‘sons’) of these crosses that inherited the construct and crossed them in turn to wild type females. We assumed that if the gene drive construct is still functional it should show a biased inheritance when the resistant target site allele had been replaced with a wild type one. Indeed in these sons we saw a significant increase in the transmission of the gene drive to their progeny, but the observed rate (average inheritance 60.13% +/-13.9%; total RFP+ progeny 59.6%) was much lower than that previously observed (~99% inheritance) (Hammond et al 2016)). A similar phenomenon of reduced homing has been observed in the offspring of another mosquito species inheriting a maternal copy of the drive construct when the same *vasa* germline promoter was utilize to transcribe the Cas9 nuclease ^7^. The reduced gene drive activity in the immediate offspring of heterozygous mothers was attributed to the persistence of maternally-deposited Cas9 in recently fertilized embryos leading to double stranded DNA breaks being repaired preferentially by end-joining mechanisms before paternal and maternal homologous chromosomes are aligned. Consistent with this explanation in the subsequent generation males (‘grandsons’) that had received a paternal copy of the gene drive had exceptionally high homing rates, with 97.5% of progeny inheriting the gene drive (Fig 3C). The drop in homing seen in sons receiving a maternal dose of Cas9 (59.6% transmission cf 97.5% grandsons) allows us to estimate an ‘embryonic end joining’ rate of 79.6% of wild type alleles being converted to cleavage-resistant alleles. This rate of embryonic end joining is much higher than that observed in the germline at or prior to meiosis (~1%, ^10^) and is predicted to reduce the rate of spread of the gene drive, due to a reduced frequency of cleavable alleles ^15^, and an increased rate at which restorative resistant alleles can arise and be selected. To investigate whether cleavage due to maternally deposited Cas9 can sometimes be followed by HDR, we note that an expected signature of embryonic homing of a resistant allele would be novel hybrid haplotypes surrounding the DSB due to partial conversion of the haplotype surrounding the wild type allele to the haplotype surrounding the resistant allele. Looking in detail at the most abundant resistant allele in each cage we failed to see such a signature and all resistant haplotypes were already pre-existing in the population (Supp Fig 1), suggesting that if this phenomenon is occurring then the resection following cleavage and resultant conversion is encompassing a section longer than the ~300bp covering our sequenced amplicon.

Our genetic analyses have shown that activity of the gene drive can lead to production of resistant alleles that fail to restore fertility, and others that substantially do restore fertility, and that the likelihood of producing these resistant alleles is higher in zygotes from mothers heterozygous for the gene drive than later in the germline. To better understand the likely effects of these processes on the spread and impact of a gene drive, we constructed a deterministic model that included two classes of resistant allele (in-frame and frame-shift, denoted R1 and R2, respectively), and having separate rates for cleavage, HDR, and EJ for maternally-derived and germline activity (see Supplementary Methods and Supp. CDF). Using this model and baseline parameter values from the single-generation crosses, we can generate the expected dynamics of allele frequencies over the 25 generations of the experiment (Fig 4). For example, the model predicts that at G12 the original wild type allele will be 9.3% of all non-drive alleles, while our observed rates were 6% and 0.4% in cage 1 and 2, respectively (Fig 2B, Supp Table 1). Importantly, for the gene drive itself, the model captures the essential aspects of the observed dynamics, showing an initial increase in frequency followed by an eventual loss, though in earlier generations our observed frequencies exceeded the predicted frequencies. To investigate what might explain this discrepancy, we examined the effect of varying each of the different parameter values in the model. In particular, small variations in the fertility of females heterozygous for the gene drive (due to partial dominance of the phenotype caused by the gene drive) had the most power in increasing match between observations and expectations. Keeping the experimental estimates of all other parameters unchanged and using a least squares regression provides the best-fit occurring at a dominance coefficient of 0.70 (Fig. 4B and Supplementary Methods), compared to our previous direct estimate of 0.9, with lower confidence limit of 0.86 ^10^. We also used our model to investigate the potential impact of HDR after cleavage caused by maternal deposition of Cas9. Perhaps surprisingly, varying this parameter has very little effect on the expected rate of resistance emergence when the rates of meiotic homing are as high as we observe.

**Figure 4.**
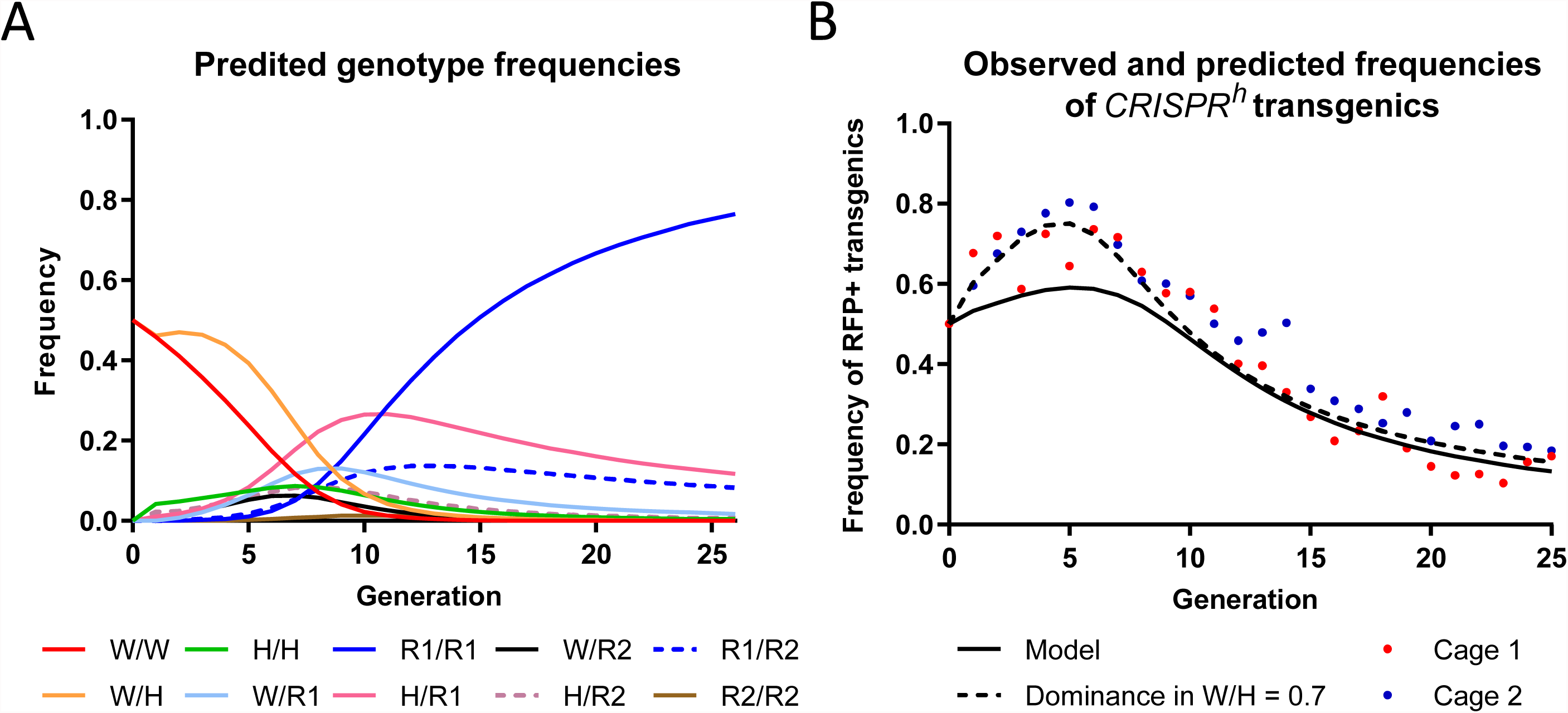
Comparison of observed data with model predicting frequencies of gene drive and resistance alleles. **(A)** Expected genotype frequencies according to the model described in the text and considering the four following target site alleles: wild type (w), CRISPR^h^ gene drive (h), resistant and in-frame (r1), resistant and out of frame (r2). We used our best experimental estimates of the considered parameters: homing rate (*e*) as 0.984, the dominance of the fertility effect due to leaky somatic expression in females heterozygous *(w/h)* for the gene drive as 0.907, meiotic end joining rate (γ_m_) as 0.01, embryonic end joining rate (γ_e_) as 0.796. **(B)** Our observed gene drive frequencies were compared against model predictions using our best experimental estimates (solid black line) and using the best-fit value (0.70 cf 0.907) for dominance of the heterozygous fertility effect in females (dashed black line).

## DISCUSSION

We have analyzed the dynamics of a gene drive deliberately designed to impose a fitness load on a population while monitoring in detail the nature of resistant or compensatory mutations, the frequency with which they arise at the target locus and the rate at which they are selected. As with any control approach aimed at suppressing an organism ‘push back’ from the target organism is to be expected. One of the advantages of the types of modular gene drives proposed here is that contingency in planning for and overcoming resistance can be foreseen and built into the system in a number of ways. For example the use of multiple gene drives targeting separate sequences has long been considered an essential pre-requisite for any gene drive intended as a functional vector control tool ^16^ and the ease with which the guide RNA expression constructs can be multiplexed lends CRISPR-based gene drives this flexibility ^9, 17^. The nuclease target site in the gene AGAP007280 described in this report was not chosen according to any prioritisation based on high levels of sequence conservation that would imply functional constraint – a feature expected to mean that resistant mutations are less likely to restore function of the gene. Clearly the choice of the target must be guided by now available extensive genomic data revealing cross-genus sequence conservation across different Anopheles species ^18^, as well as by a detailed knowledge of the natural occurring variation currently existing in *Anopheles gambiae* target populations as revealed by resequencing of large numbers of wild-caught mosquitoes ^19^. This data revealed *a posteriori* that for the target site in AGAP007280 used here there is pre-existing variation in at least 8 of the 20 nucleotides covered by the gRNA. Going forward low tolerance of sequence variation at the target site should be a key criterion for designing a gene drive. This is all the more important given that that one of the key drivers in the generation of resistant alleles is the nuclease activity itself and the end-joining that can mediate its repair. Our results show that a significant proportion of these alleles are created as a result of maternally deposited nuclease in the early zygote where end-joining repair massively predominates over homology-directed repair. This maternal effect is likely to be simply solved either through the use of more tightly regulated promoters to restrict nuclease expression to the early germline or through the addition of destabilising modifications to the nuclease, either of which are expected to reduce the perdurance of the gene drive component in the embryo. Nonetheless, even the low rate of end-joining that we observed during gametogenesis alone is likely to quickly compromise a gene drive relying on a single target site, as our model and that of others shows ^11, 17^, unless the requirements for targeting multiple functionally constrained sites are met. An additional consideration in the choice of target site may take into account the propensity for a particular double strand break to be repaired more readily into a resistant, restorative (r2) allele, for example due to microhomology either side of the cleavage site that more readily re-creates an in-frame allele than a frameshift allele.

Our approach of pooled sequencing of a targeted region allowed us to reliably detect even low frequency signatures of gene drive activity and reveal the complex dynamics of different genotypes emerging over time. Certainly for the future improvement of gene drives it will be important to have a faster method to triage for the most robust gene drives least prone to resistance without a multi-generational cage experiment, a laborious and time consuming process that should be reserved for more extensive evaluation of the best candidates. A simple way to do this would be to apply the method of amplicon sequencing described here in a screen where all generated mutant alleles are balanced against a known null allele to see if they restore function to the target gene.

The potential for rapid emergence and spread of resistance highlights on the one hand some of the technical challenges associated with developing a gene drive while on the other hand highlights how the option of intentionally releasing a resistant allele in to a population could be a simple and effective means of reversing the effects of a gene drive.

## METHODS

### Gene drive generational cage experiments

These experiments were essentially as described before in Hammond et al. (2016). Briefly, in the starting generation (G_0_) L1 mosquito larvae heterozygous for the CRISPRh allele at AGAP007280 were mixed within 12 hours of eclosion with an equal number of age-matched wild-type larvae in rearing trays at a density of 200 per tray (in approx. 1L rearing water). The mixed population was used to seed two starting cages with 600 adult mosquitoes each. For subsequent generations, each cage was fed after 5 days of mating, and an egg bowl placed in the cage 48h post bloodmeal to allow overnight oviposition. After allowing full eclosion a random sample of offspring were scored under fluorescence microscopy for the presence or absence of the RFP-linked *CRISPR^h^* allele, then reared together in the same trays and 600 were used to populate the next generation. After a generation had been allowed the opportunity to oviposit, a minimum of 240 adults were removed and stored frozen for subsequent DNA analysis.

### PCR of Target Site and Deep Sequencing library preparation

For the sequence analysis, a minimum of 240 adult mosquitoes taken at generation G_2_ and G_12_ of the cage trial experiments were pooled and extracted en masse using the Wizard Genomic DNA purification kit (Promega). A 332 bp locus containing the target site was amplified from 40 ng of genomic material from each pooled sample using the KAPA HiFi HotStart Ready Mix PCR kit (Kapa Biosystems), in 50 μl reactions. Specially designed primers that carried the Illumina Nextera Transposase Adapters (underlined), 7280-lllumina-F <u>(TCGTCGGCAGCGTCAGATGTGTATAAGAGACAG</u>GGAGAAGGTAAATGCGCCAC) and 7280-lllumina-R <u>(GTCTCGTGGGCTCGGAGATGTGTATAAGAGACAG</u>GCGCTTCTACACTCGCTTCT) were used to tag the amplicon for subsequent library preparation and sequencing. The annealing temperature and time were adjusted to 68°C for 20 seconds to achieve the minimum off-target amplification. In order to maintain the proportion of the reads corresponding to particular alleles at the target site, the PCR reactions were performed under non-saturating conditions and thus they were allowed to run for 20 cycles before 25 μl were removed and stored at -20°C. The remnant 25 μl were run for an additional 20 cycles and used to verify the amplification on an agarose gel. The non-saturated samples were used to prepare libraries according to the Illumina 16S Metagenomic Sequencing Library Preparation protocol (Part # 15044223 Rev.A). Amplicons were then purified with AMPure XP beads (Beckman Coulter) followed by a second PCR amplification step with dual indices and Illumina sequencing adapters using the Nextera XT Index Kit.

After PCR clean-up via AMPure XP beads and validation performed with Agilent Bioanalyzer 2100, the normalized libraries were pooled and loaded at a concentration of 10 pM on Illumina Nano flowcell v2 and sequenced using the Illumina MiSeq instrument with a 2x250bp paired end run.

### Deep Sequencing analysis

Sequencing data of the amplified genomic region were analysed using available tools and developed scripts in R v3.3.1. Raw reads were cleaned up for low quality and trimmed for the presence of adapters using Trimmomatic v.0.36 ^20^.

Paired end reads were merged together in order to reconstruct the whole amplicon sequence using PEAR v0.9.10 ^21^. Resulting assembled identical fragments were then clustered using fastx_collapser module from FASTX v0.0.13 suite (<u>http://hannonlab.cshl.edu/fastx_toolkit/</u>) and aligned to the reference amplicon with vsearch tool v2.0.3^22^ which implements a global alignment based on the full dynamic programming Needleman-Wunsch algorithm. We considered for downstream analysis only sequences represented by at least 100 reads in each dataset. The .blast6 output files from the alignment phase were parsed by ad hoc written R scripts to identify sequence variants containing insertions and/or deletions in the target site. The quantification of each allelic variant was measured as relative alternative allele frequency by summing up the reads representing that particular variant in the dataset. Finally, for each identified variant, we examined the single nucleotide variants (SNVs) along the full amplicon and selected the ones with a minimum alternative allele frequency of 2.5% for the purposes of haplotype calling.

### Modelling

We use a Wright-Fisher type model to explore how the dynamics of gene-frequencies depend on underlying parameters. We suppose there are four possible alleles in the population at any time: Wildtype (W), driver allele (H), and two types of mutant allele that are resistant to homing, R_1 which is fully functional and R_2 which is recessive but non-functional (i.e. R2/R2 type individuals are sterile). Given that we failed to observe any obvious reduction in any of the 5 classes of in-frame alleles that had been selected in our cage experiment we assumed no fitness cost of the R1 allele, though our model permits this to vary. We assume allele pairs segregate at meiosis according to Mendelian inheritance except for W/H zygotes where segregation may be distorted by cleavage followed by either homing or non-homologous repair. Our model also allows the possibility that eggs from females with at least one H allele will contain the driver nuclease (regardless of the eggs own gamete type), in which case cleavage and repair may occur in the embryo. The mathematical details of the model are given in the Supplementary Methods. We incorporated estimates of the parameters expected to affect the rate of spread of the gene drive and the generation of any resistant alleles over time, into a deterministic model in computable document format (.cdf) (Wolfram CDF player, free to download at <u>https://www.wolfram.com/cdf-player/</u>) that allows the user to change each parameter and to predict the relative frequencies of different genotypes while comparing against our observed data (see Supplementary Methods and Supplementary Files). We incorporated our experimental estimates of homing rate (e) as 0.984, the dominance of the fertility effect due to leaky somatic expression in females heterozygous ( *w/h)* for the gene drive as 0.907, meiotic end joining rate (γ_m_) as 0.01, embryonic end joining rate (γ_e_) as 0.796.

### Individual Assays of Mosquito Fertility and Transmission of Gene Drive

Individual females containing at least one copy of the RFP-linked *CRISPR^h^* gene drive were selected from the G_20_ generation and allowed to mate with 5 wild type male mosquitoes, essentially as in Hammond et al.^10^. The fecundity of females and transmission of the gene drive was measured by counting larval offspring positive for the RFP marker as a proxy to assess the frequency of gene drive transmission. To check mating status of females, spermatheca were dissected and examined for the presence of sperm. Unmated females were censored from the fertility assay.

Sons of each gene drive mother from the G_20_ generation were kept together and allowed to mate in groups of approximately 5 males with an equal number of wild type females and assessed for rates of transmission of the gene drive. The male offspring of these sons (grandsons) inheriting the drive from their fathers were in turn assessed in the same way, keeping lineages separate.

## Acknowledgements

We are grateful to M. Astrakhan for technical assistance, to N. Windbichler, P. Papathanos for useful discussion and to A. Burt for helpful feedback. This work was supported by a grant from the Foundation for the National Institutes of Health through the Vector-Based Control of Transmission: Discovery Research (VCTR) program of the Grand Challenges in Global Health initiative of the Bill & Melinda Gates Foundation

---

**Supplementary Figure 1.**
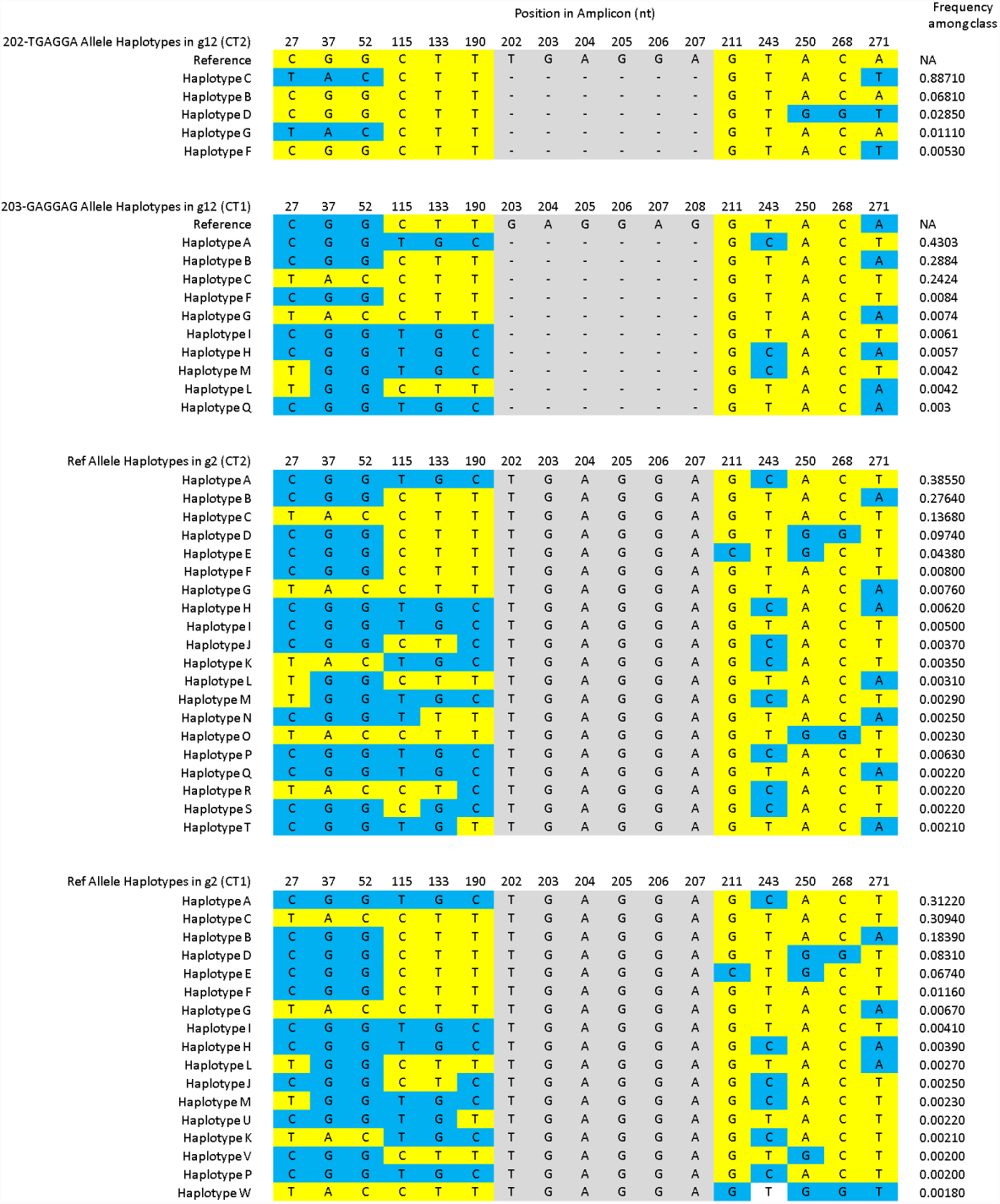
Resistant target site alleles showing selection are formed independently on a wide range of haplotype backgrounds. The presence of polymorphic SNPs surrounding the target site and circulating at various frequencies in the laboratory wild type colony allowed us the resolution to identify a variety of haplotypes on which target site indels may have been formed. The most prominent target site indel in each cage replicate in the G_12_ generation was analysed and the number of haplotypes containing the respective indel and the frequency of each haplotype was calculated. A measure of the diversity of pre-existing reference haplotypes present in the colony was obtained by examining the nature and frequency of haplotypes surrounding the wild type target site allele in the early G_2_ generation. In both replicates there were no unique haplotypes containing the indel that were not already pre-existing in the starting population. The relative frequency of haplotypes surrounding a given target site allele are also displayed.

**Supplementary Table 1 Target site allele frequencies at g2 and g12 generations**

**Supplemetary Table 2 Gene drive transmission in offspring of *CRISPR^h^/R* receiving a maternal copy of gene drive, and subsequent paternal copies**

